# Simultaneous fluorescence imaging of tilted focal planes at two depths in thick neural tissue: Implementation with remote focus in a widefield electrophysiological microscope

**DOI:** 10.1101/839050

**Authors:** Bruno Lagarde, Noah Russell, Elric Esposito, Laura Desban, Claire Wyart, David Ogden

## Abstract

Wide-field imaging conventionally results in a single image plane oriented perpendicular to the optical axis. However, in brain slice or *in vivo* recording, neuronal or circuit morphologies lie in arbitrarily tilted planes. Consequently the spatiotemporal advantages of wide-field non-scanned imaging are lost because of the time required for stepwise focal readjustments to view an entire neuron or network. We describe an application of remote focus that views simultaneously two planes separated by up to 100 µm, each with variable tilt from the conventional image plane. This permits fluorescence detection of ion fluxes or membrane potential across neuronal compartments and their correlation with electrical activity. Further, two fluorophores can be viewed simultaneously in each plane.

We show (i) neuronal images tilted to optimise simultaneous aquisition of somatic, dendritic and axonal compartments; (ii) networks viewed simultaneously at 2 depths separated by up to 100 µm, (iii) widefield imaging at 30 Hz of Gcamp5 fluorescence during spontaneous spiking in motoneuron layers of zebrafish spinal cord separated by 30-40 microns.

## Introduction

Electrophysiological recording in neuroscience is often combined with fluorescence microscopy of indicators of cytosolic ion concentrations or membrane potential, imaged with EMCCD or sCMOS cameras. This generally has the advantage over scanning of simultaneous time-resolved data acquisition at high speeds and widefield exposure. With modern cameras frame rates of 100 Hz can be obtained at sub-µm resolution over a 200 µm field. The fluorescent indicators are usually perfused into the cell from a whole cell recording pipette which also records spike output or synaptic current; alternatively, indicators may be fluorescent proteins transfected by transgene or viral methods and electrical recording with extracellular probes. Experiments can correlate the high time resolution of electrical recording with compartmentalized detection of ion concentrations and fluxes, or image local membrane potential (for a review of methods see Canepari et al., 2013). A technical problem in brain slice recording is that neurons survive if their neurites have not been damaged, consequently surviving cells recorded near the slice surface lie mainly outside the conventional focal plane normal to the optical axis. Similarly, neurons or networks *in vivo* have orientations constrained by points of access in the skull and are rarely in the conventional focal plane. Consequently the advantages of full-field non-scanned imaging are lost in re-focusing, considerably slowing data aquisition. Here we describe practical implementations of remote focus (Botcherby et al., 2008) adapted to suit electrophysiological experiments in conventional commercial microscopes. These permit the focal plane to be tilted in an arbitrary orientation and translated axially to coincide with the geometry of neurons, glial cells or networks of interest within the slice. In the simplest implementation time-resolved local changes of fluorescence can be recorded simultaneously over the major plane of the neuron or network, evoked by presynaptic electrical stimulation, by wide-field or localized excitation or inhibition with uncaging or optogenetic techniques. Further, by separating the two images by polarisation and projecting onto adjacent regions of the same camera sensor, or separately on two cameras, simultaneous optical recordings can be made at two depths separated by 100 µm without introducing additional aberrations or changing the magnification. By incorporating a second remote focus, images at two depths and different orientations can be viewed. Furthermore, since the images are separated by polarisation they can be further separated to view two colours in the emitted fluorescence. In the applications described here the primary aim is to improve time resolution of fluorescence imaging in thick specimens by removing the need to refocus, and to be able to image simultaneously at two planes on the z-axis

The remote focus method was introduced and described in several configurations by Botcherby et al. (2008). It uses the principle, due to Maxwell (1858), that an image with optical path lengths equal to those within the specimen can be viewed without spherical aberration (see also Born and Wolf, 2002). Maxwell showed that the magnification achieving this condition is the ratio of refractive indices of media in the specimen and remote image, since these multiply the geometric path lengths. Conventional microscopes which magnify directly do not satisfy this condition and tilted planes or multiple planes at wide separations cannot be viewed without spherical aberration. To generate a remote image with minimal aberration the correct magnification is achieved by demagnifying with a second objective to form an image with magnification M1/M2=n1/n2, the ratio of refractive indices at the specimen and image. The different planes in the remote image are reflected with a moveable mirror and separated from the primary image by polarisation. The general arrangement is presented in the schematic of Fig1A, which shows the primary imaging objective O1, the remote objective O2, the tube lenses and the polarization splitter that separates images from the two paths before projection to the detectors. Focal planes in the remote image space of O2 are sampled by the mirror and reflected to the camera. For slice or *in vivo* recording primary objectives O1 are usually water dipping, combining with an air objective usually used as the remote objective at O2 requires a ratio of magnifications M1/M2 = n1/n2 = 1.33. The final magnification in the viewed image is determined by the projection lens to the camera. The simple scheme with one remote focus objective is illustrated in Fig.2B; the arrangement with two remote paths, permitting two tilted images at different depths, shown in Fig 4.

**Fig.1.**
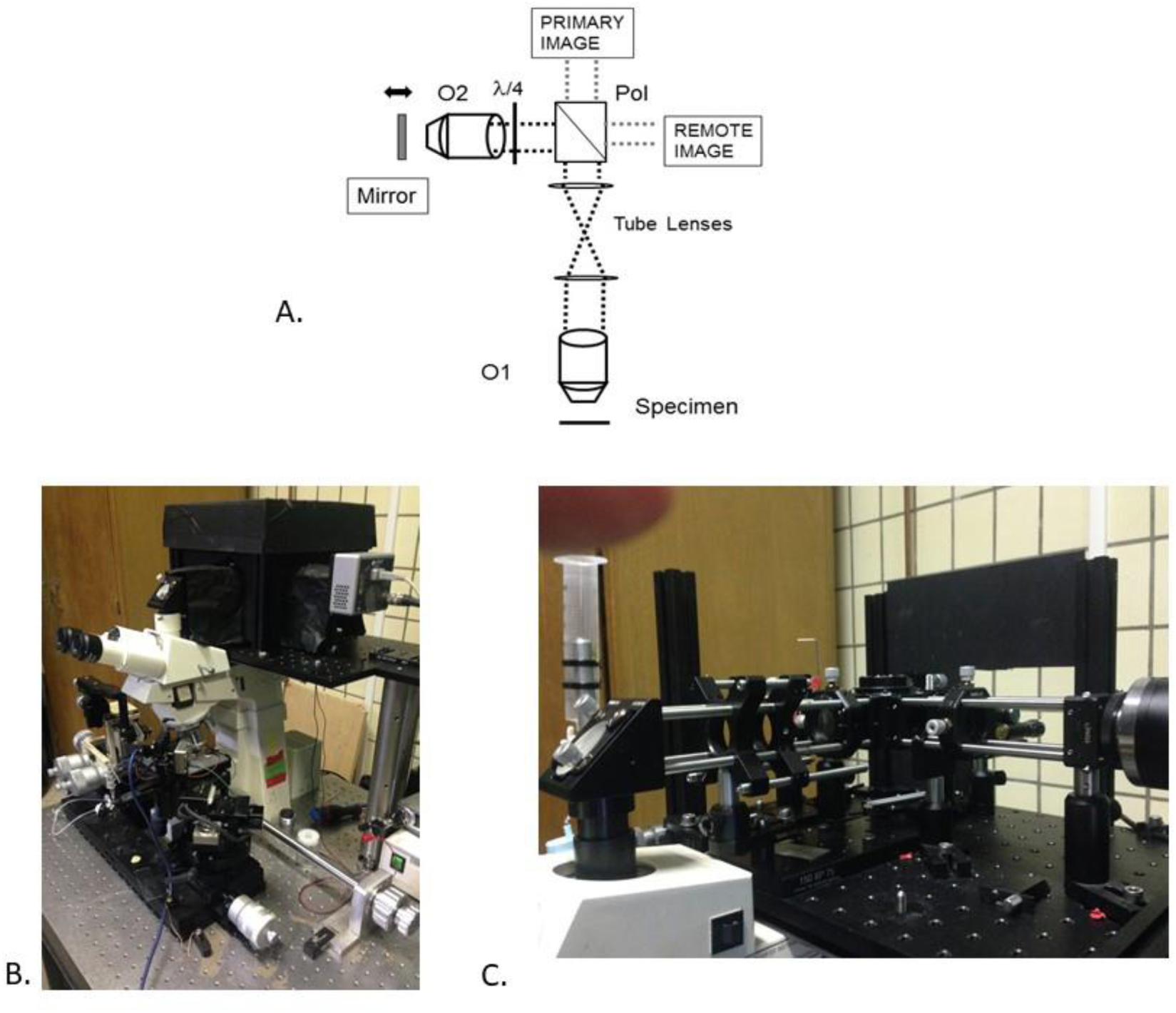
A. Outline of the remote focus optics. Primary objective O1 forms an intermediate image projected by first tube lens and viewed by the second tube lens (of O2). One polarisation is passed through the polariser cube and can be projected to form the primary image (as seen by the oculars). The orthogonal polarisation is reflected to remote objective O2, the image reflected from a translatable gimbal mirror. The 2 passes of the λ/4 plate change polarisation by λ/2; the image is transmitted through the polarising cube and projected to a camera as the remote image. In Fig 2 schematic, the primary image is also retarded λ/2 by reflection through a λ/4 and transmitted to the camera as a reference image useful for alignment. B, C Arrangement of remote focus components on a conventional slice microscope, here a Zeiss Axioskop FS1. 1B (Left panel) experiment setup with blackout card supported on grooved pillars. 1C. (Right panel) showing the 30 mm cage system components and 30/60 mm cage system support projecting the microscope intermediate image to the remote focus optics.

**Fig.2.**
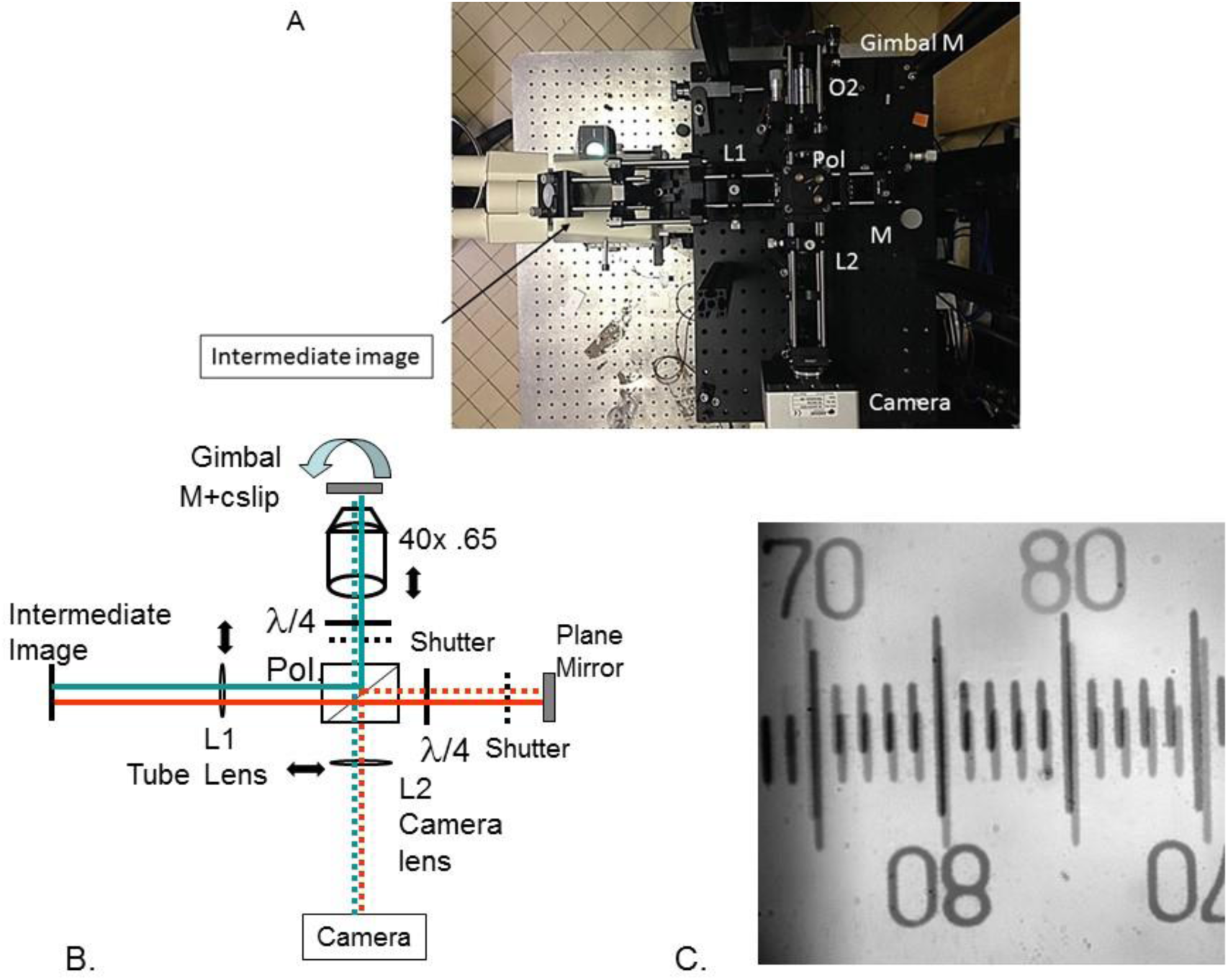
A. Photograph of the breadboard with 30 mm /60 mm cage components. Position of intermediate image and key components indicated: L1-tube lens; Pol-polarising beam splitter; L2-camera lens; Gimbal Mirror; M-plane mirror. O2-remote objective. Fig 2B. Schematic of the optical components. Tube lens L1 (170 mm or 120 mm; see text), camera lens L2 (150 mm or 100 mm), camera Andor Luca or Ixon (EMCCD, 1000×1000 or 512×512, 12 mm sensor). Red path is non-remote reference path; blue path is through the remote objective O2. Fig 2C. The primary and remote images of a stage micrometer (10 µm /div) viewed simultaneously from both paths to facilitate rotation and tilt alignment of the polarizing beam splitter (Pol). Note image inversion in the remote path.

The optical performance of remote focus has been assessed by Botcherby et al. (2008), by Anselmi et al. (2013) and by Qui et al. (2014). These studies show that spherical aberration in the remote image is negligeable up to depths of 150 µm from the primary focus. Viewing different planes reflected from the remote image is most efficient by selecting orthogonal polarizations with a beam splitting cube and quarter wave retarders as shown in Fig 1A, Fig 2 and Fig 4. The polarizing cube is placed in the infinity space between the remote objective and its tube lens, which collimates light from the intermediate image of the primary objective. The pupils of the 2 objectives are imaged together by the tube lens combination preferably in a 4f configuration, and to maintain resolution the cone angle of the remote objective to the optical axis should be greater than that of the imaging objective (Botcherby et al., 2008). Measurements by Anselmi et al.,(2011) of the Strehl ratio in a similar configuration, with 40x 0.8 NA water dipping primary objective and 40x 0.65 NA air remote objective, showed no additional aberrations were introduced with z-displacements of +/− 150 µm from the primary focus. A maximum tilt angle of 14° from the conventional focal plane was limited mechanically by the working distance of the remote objective O2.

Methods to view two or more widely separated focal planes with a single imaging objective without remote focus have not been reported. In studies in which spatial definition rather than time resolution was the primary aim the z-range achieved was limited to 1-2 µm by the need for large compensatory axial excursions of the camera (see for example Prabhat et al., 2004). Unlike wide field remote focus they do not have 100 µm z-range nor the simplicity of implementation with a single imaging objective. Light sheet illumination requires an additional objective near the specimen, limiting their use in electrophysiological applications requiring micromanipulation.

Recent developments of the remote focus method have been in laser scanned imaging where the fast z-scan capability of a small mirror provides 3D scanning in two-photon microscopy. However, time-resolved imaging combined with electrophysiology is more readily applied in wide-field camera configurations. The aims here were (i) to adapt the optical train to an upright ‘slice’ microscope, permitting simultaneous electrophysiological measurements, (ii) to simplify alignment by providing a simultaneous reference image to the camera from the non-remote polarization, and (iii) to demonstrate the ability to view two widely separated depths in the tissue simultaneously by viewing the two orthogonal polarisations, either with an image splitter or on separate cameras.

## Methods and alignment

The design makes use of the space behind the trinocular head of upright microscopes to mount the remote system on a breadboard supported above and behind the area used by micromanipulators or for stage translation. Fig.1 B and C show the arrangement on a Zeiss Axioskop FS1 microscope with a conventional trinocular camera port viewing the intermediate image from the tube lens of the microscope. The additional optics for the remote focus are assembled in a 30 mm cage-rod system on a breadboard supported on pillars. The Zeiss Axioskop FS1 is head-focusing, to permit vertical translation when focusing the intermediate image was projected into a 90 deg mirror holder with elliptical mirror and a mechanically isolated tube suspended within the camera port. The intermediate image formed by the 165 mm tube lens of the Zeiss Axioskop appears at the exit of the right angle mirror holder indicated in Fig.1B. The 30/60 mm cage system permitted repositioning of the microscope when occasionally required for example changing O1.

A schematic of the external optical path is shown in Fig. 2A. The intermediate image is viewed by lens L1 (the tube lens of the remote objective) and projects to a polarising beam splitter with rotational and tilt adjustment. One polarisation passes through to an arm comprising a λ/4 retarder in a rotation mount and a plane mirror. Light reflected from the mirror is rotated λ/2 by 2 passes through the λ/4 retarder and is reflected in the polarising cube to the camera arm. After projection this image corresponds to that seen in the oculars. The second polarisation is reflected into the perpendicular path, sequentially through λ/4 retarder in a rotation mount, a simple shutter, the remote objective O2 and is reflected from a plane mirror in a tilting gimbal mount. The remote image reflected from the gimbal mirror has polarization changed by λ/2 and passes the polarising cube to the camera arm. Thus both polarisations pass to the camera arm, one directly from the intermediate image, the other via the remote objective, with the same magnification. Transmission efficiency to the camera is maximized by rotation of the λ/4 retarders.

### Calculation of the lens parameters

For single neuron recording the objective O1 fitted to the Zeiss was a 63x 0.9NA Leica water dipping objective (requiring an rms-25 mm adaptor). The Leica objective is designed for a tube lens of 200 mm focal length, when using the Zeiss tube lens of 165 mm the magnification at the intermediate image plane is 52x. The remote objective O2 was a 40x 0.65 NA Olympus air objective with 0.17mm cover glass correction. The condition that the magnification ratio should be the same as the ratio of refractive indices was adjusted by the tube lens, L1, of O2. For the Olympus objective O2 the magnification is 40x for 180 mm tube lens, consequently to achieve a ratio of M1/M2 = n1/n2 = 1.33, L1 is calculated as 175 mm. An AR coated lens of 170 mm was used, positioned 170 mm from both the pupil of O2 *via* the polarizing cube and from the mirror in the reference arm. The position of the breadboard was adjusted with respect to the microscope such that L1 was 170 mm from the intermediate image plane located at the exit of the 90 deg reflector mounted in the trinocular head. A camera with 12 mm sensor was used (Andor Luca EMCCD, 1000×1000 pixels, or Andor Ixon 887, 512×512) and a lens L2 of 150 mm focal length projected the image to the camera. This configuration resulted in a magnification of 1.38x from sample to remote image and a total magnification of 46x from sample to camera.

The remote objective O2 was mounted in a Linos micrometer stage (Z-Fine adjustment M, Quioptic, resolution 1 µm) supported on the 30 mm cage/rod system. The gimbal mirror mount (Thorlabs) was post-mounted on the breadboard. A 1 inch protected silver mirror was used with No 1.5 coverglass glued with index matched adhesive to compensate the cover glass correction in O2.

For experiments shown in Fig 8 requiring a larger field the primary objective was changed to a Leica 40x 0.8 NA water objective at O1 with O2 unchanged, and a 120 mm lens used at L1 to reduce the magnification correspondingly in the remote path (calculated value 111 mm for magnification ratio 1.33); the total magnification was 29x at the camera. The position of the microscope relative to remote objective was readjusted by moving the platform and utilizing the 30-60 mm cage assembly to adjust the position of L1 relative to the intermediate image plane. In this case L1 was 120 mm from the intermediate image and from the pupil of O2. For Fig 9, the primary objective was changed to Olympus 40x 0.8w (corrected for 180 mm tube lens) L1 changed to 125 mm to give n1/n2=1.33, and the camera lens L2 to 150 mm giving a 180 µm field at the camera.

### Alignment procedure

The steps in alignment are explained with reference to Fig 2. The resolution is determined by choice of O1 and O2 and the lens L2 determines the magnification at the camera sensor. The tube lens L1 is calculated as shown above. The steps in alignment are the following: (i) The camera is used in alignment by setting the image to infinity. To do this here the 90-degree adaptor and L1 are removed by disconnecting the rods supporting the adaptor. The position of projection lens L2 adjusted for best focus on a distant target and locked in position. (ii) the position of lens L1 is adjusted to be 1 focal length from the pupil of O2 and from the plane mirror in the non-remote arm *via* the polarising cube (iii) The microscope and breadboard are positioned so that the intermediate image (at the exit of the 90 degree adaptor here) is 1 focal length from L1. (iv) The microscope intermediate image of a stage micrometer is initially viewed with the oculars focusing O1 and the output switched to the remote focus/camera path. (v) To align the polarising cube a cube/rod system alignment tool with central 1 mm aperture is suspended on the cage rods at the intermediate plane and illuminated with brightfield light *via* the microscope. L1 and L2 are centred in their housings. Plane mirrors are inserted in the shutter slots in each arm (13 mm mirrors in Thorlabs shutters) and the rotation and tilt of the cube adjusted to align the 2 light spots with each other centrally on the camera; additional centring of coincident spots on the camera by lateral adjustment of L2 and L1. After removing the alignment tool, the image of a stage micrometer with brightfield illumination was used to check the field size and registration of images simultaneously from the 2 paths (Fig 2C). (vi) The remote focus gimbal mirror was aligned with the optical axis and the reference position locked; this was facilitated by a 30 mm cage system plate positioned on extensions of the upper cage rods above O2, providing a mechanical reference normal to the optical axis. After removing the mirror/shutter from the remote path the remote image was set up to be the same as the direct image by fine focus Z-adjustment of the remote objective O2 position, by rotation of the λ/4 retarder to maximise throughput, and by adjustment of the gimbal controls to give the same image as seen in the non-remote path (note image inversion in the remote path). Both images were projected to the camera and aligned further if needed by rotation/tilt of the polarising cube. Fine adjustments to the z-position of L1 were made to give the same magnification of both images of the stage micrometer at the camera. The position of images compared to the ocular image was by adjusting tilt of the mirror at the trinocular head and x-y translation of L1 and L2.

### Zebrafish care and strains

*Danio rerio* embryos of the AB and Tubingen Longfin (TL) genetic backgrounds were raised in egg water, at 28.5°C until 75% epiboly and then at 26°C to delay development for experimentation until the 24-somite stage. GCamP5-G-positive embryos originating from the cross of *Tg(mnx1:GCamP5G)* (Sternberg et al., 2018) with wild type adults were dechorionated, mounted laterally in 1.5% agar at the 24-somite stage, anesthetized transiently using MS-222 (Sigma-Aldrich, USA) and paralyzed with injections of 0.5 µM α-bungarotoxin (Sigma-Aldrich, USA) into caudal muscle of the tail. Experiments were performed at the 26–30 somite stages at room temperature (23°C). All procedures on zebrafish embryos were approved by the Institutional Ethics Committee “Charles Darwin” at the Institut du Cerveau et de la Moelle épinière (ICM), Paris, France, and received subsequent approval from the EEC (2010/63/EU).

## Results

### Optical parameters

#### Efficiency of Transmission in remote focus

The fraction of light transmitted through the remote focus paths from the intermediate image was tested at different points with a 1 cm square photodiode (Thorlabs FDS1010 with I/V amplifier) glued to a cage/rod alignment tool. More than 50% loss is expected in each path from the intermediate image to the remote image due mainly to polarisation selection and losses due to reflection. Transmission from the intermediate image plane to the remote focus image at the camera was measured at different points with a large area photodiode in the green spectral region (510-550 nm), viewing the brightfield image of a 20 µm pinhole in the specimen focus of O1. The fraction of light transmitted from the intermediate image to the camera imaging plane *via* the remote focus path was 30%, compared with 50% expected for polarisation loss alone. The additional losses resulted mostly from the 2 passes in the remote objective; losses by reflection at the polarising cube and quarter wave retarder were found to be small. An asymmetry was found in transmission in the polarising cube. With 2 remote paths combined (scheme in Fig.5B) the throughput from the intermediate image to the camera totalled 65% in both paths.

#### Field of view

Since the magnification after the remote objective is the same in both paths to satisfy Maxwell’s criterion (Maxwell, 1858) the field size seen at the camera depends on the camera projection lens L2 and on the sensor size, 12 mm used here. The field size with a 150 mm lens at L2 was 180 µm, seen in the images of a stage micrometer of Fig 2. To image networks of interneurons (Fig8) the field size was increased by changing projection lens L2 to 100 mm focal length giving field size 320 µm diameter at the 12 mm sensor.

##### Measurement of full projected field size

To test the field size without limitation by the sensor a brightfield image of a stage micrometer was projected to a screen in place of the camera sensor. This showed that the image in the primary objective (O1 40x 0.8w Olympus) was 500 µm in the primary polarisation, whereas the orthogonal polarisation passed through the remote objective (O2 Olympus 40x 0.65 air) had field reduced to 360 µm at the same focal plane, limited by an aperture in the remote objective; an objective at O2 specified for a larger field should permit large fields of view in the remote image. In the present experiments field size was limited by the size of the camera sensor.

##### Depth and tilt test

Test slides of fluorescent beads distributed at different depths were made with 5 µm or 0.28 µm Fluoresbrite beads (Polysciences Inc) suspended in Sylgard 184, dropped onto cover glass and cured at 60 degree for 5 min. Measurements of depth and tilt were determined from the focus of the Zeiss Axioskop microscope, calibration was verified as 2 µm per division. The depths of pairs of beads at extremes of the field were measured with the Zeiss Axioskop focus in the non-remote path and brought into view, as closely as possible in the same plane, with the remote focus z-translation and gimbal mirror. As reported previously (Anselmi et al., 2011) the maximum tilt achievable with an Olympus 40x 0.65NA (wd 0.6 mm) at O2 was 14°, limited by mechanical interaction between the cover glass glued to the gimballed mirror and the objective rim. Anselmi et al (2013) reported that misalignment of the objective pupils resulting from tilt did not impair resolution.

#### Resolution in the remote image

Measurements of the resolution in the remote image compared with the primary image were made by Anselmi et al., (2011) who compared the Strehl ratios measured in the two images. They reported no additional loss of resolution over +/− 150 µm in the remote image. The point spread function of diffraction limited laser illumination at different depths and lateral positions in the remote relative to the primary image were also reported to show no distortion over a range of 180 µm (Botcherby et al 2008). Here images were taken of test preparations, refocusing in the remote path at depths 50 µm or 100 µm below the primary focus. As above, Fluoresbrite beads of 5 µm or 0.28 µm diameter suspended in Sylgard 142 to produce a 3D distribution of fluorescent beads imaged at different depths with primary and remote objectives. They were separated for display with a polarising beam splitter (Optosplit, Cairn Research) projecting the images to adjacent regions of the camera sensor (see Fig.4). Fig 3 A shows 2 beads of 5 µm diameter separated in z by 100 µm imaged simultaneously in primary (upper) and remote (lower) fields. Each is shown zoomed in Fig. 3B. Sub-resolution 0.28 µm beads in Sylgard are shown in Fig 3C at 3 depths with high zoom, showing that images viewed at widely separated depths in the remote image (50 and 100 µm) have spatial definition similar to the primary image. The images obtained in brightfield of a stage micrometer and in epifluorescence of 5 µm diameter fluoresbrite beads at zero and 100 µm separation on the optical axis are shown in Fig 4 D, E. Fig 6 shows brightfield and fluorescence images of a pollen grain, which show the resolution was maintained in the brightfield image at 100 µm defocus, however in epifluorescence the resolution appears degraded at 100 µm but not at 50 µm.

**Fig. 3.**
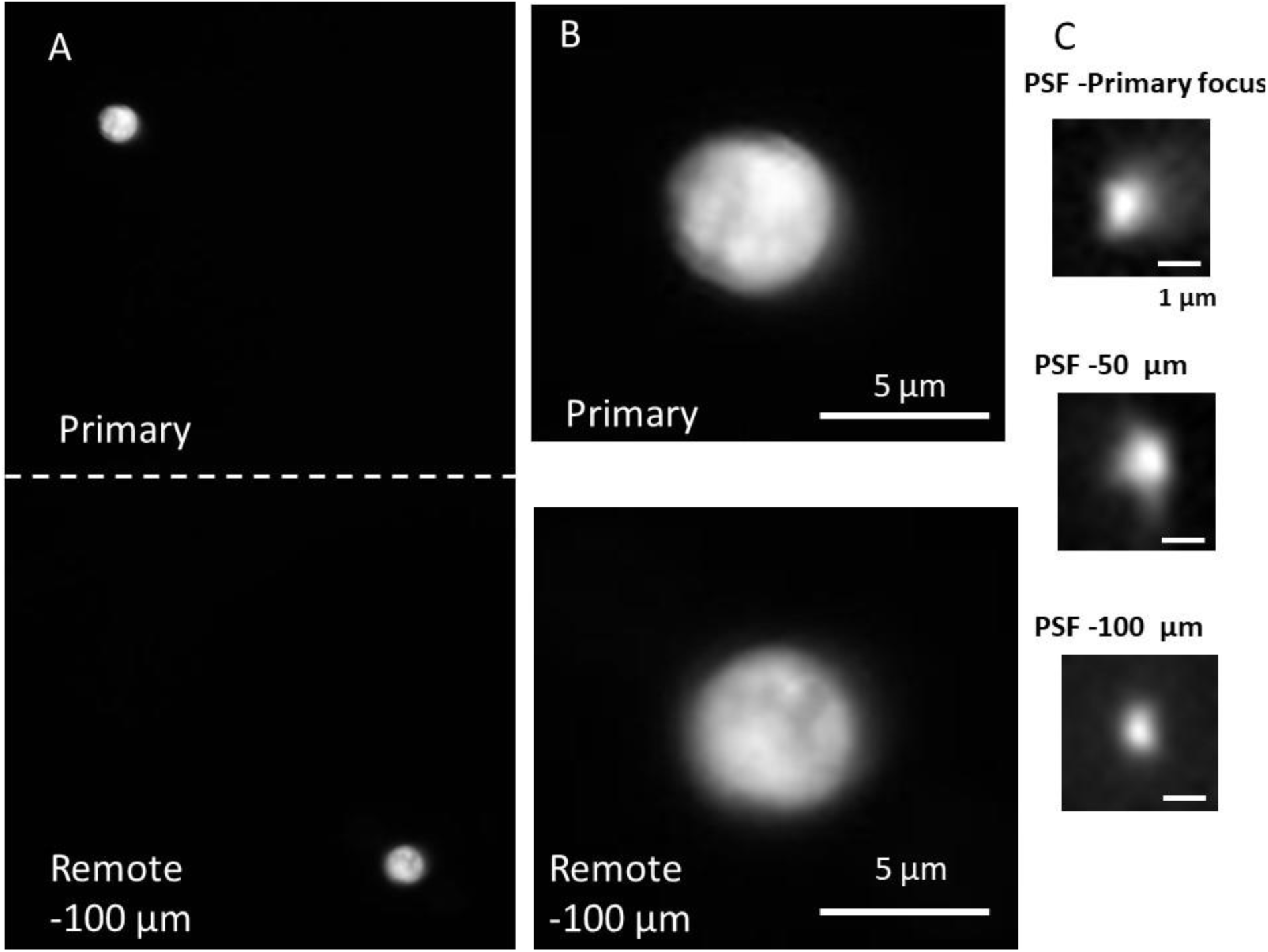
Resolution of primary and remote focus compared. A, B: Fluoresbrite beads 5 µm diameter suspended in Sylgard resin imaged simultaneously onto a 12 mm emccd sensor (exc 470/40 nm, em 525/40 nm) viewed *via* polarising Optosplit (Cairn Research). C: 0.28 µm beads suspended in Sylgard and imaged as single beads to compare PSF at primary focus (top) at defocus −50 µm in remote focus (middle) and −100 µm (bottom) in the remote image.

**Fig. 4.**
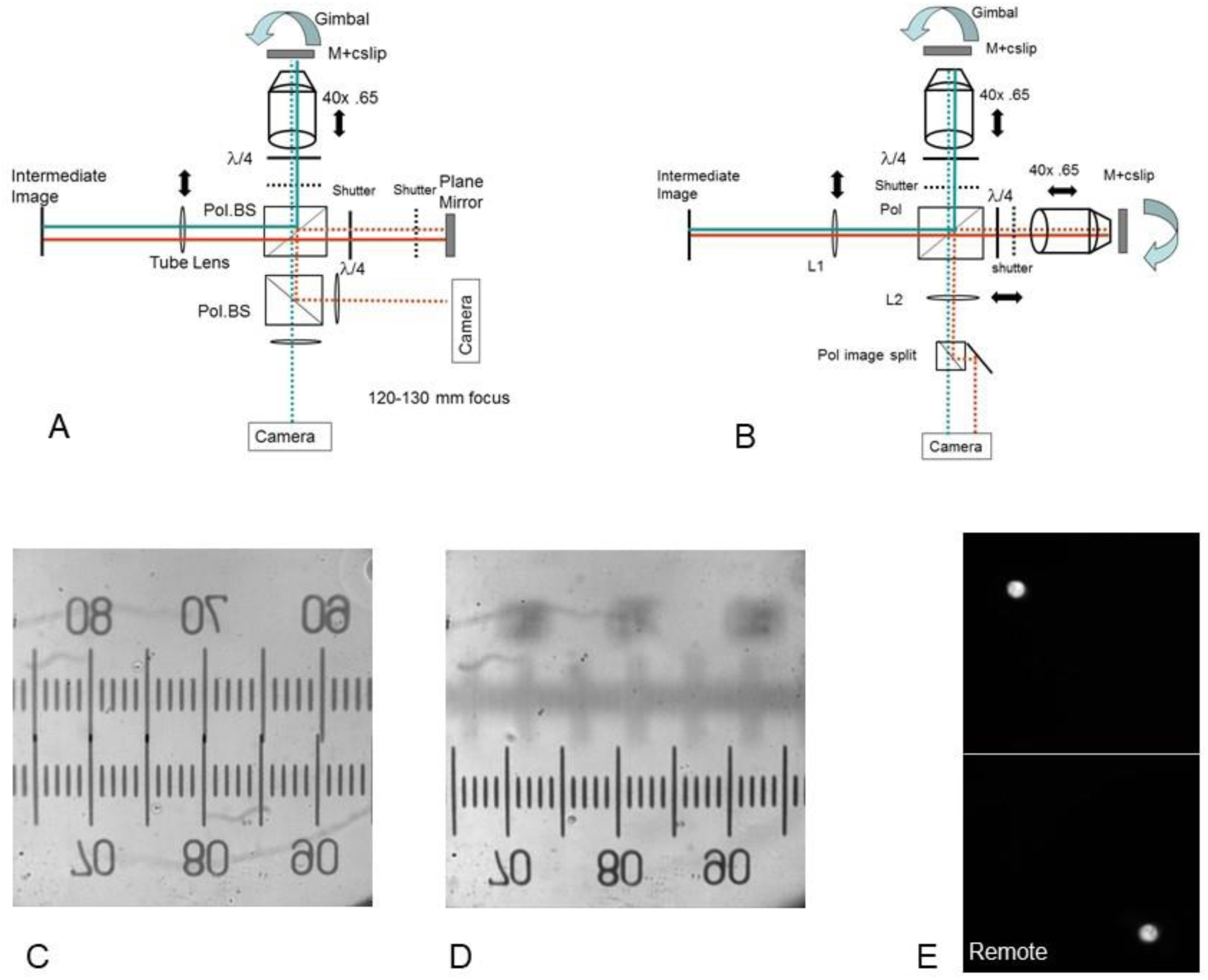
Schematic of the optical arrangement for a single or 2 remote paths to view planes at different depths in the sample. An additional polarising beam splitter projects the images with orthogonal polarisations from the 2 paths on to (A) separate cameras or (B) in conjunction with an image splitter (polarising Optosplit or Multisplit, Cairn Research) to the same camera chip. Projection lens L2 was 100 mm FL giving field size 320 µm. With a single remote path (panel 4A) the microscope primary image (O1) can be viewed simultaneously with a remote image (O2) at different depth and orientation (scheme Panel 4A); With 2 remote paths (Panel 4B) both can be at different depths from the primary focus and tilted to optimize orientation of both images independently. Panel 4C. stage micrometer (10 µm /div) viewed through direct and remote focus paths simultaneously at the same focus. Panel 4D: After displacement of the primary focus O1 by 100 µm, generating the unfocused image, the image was restored by adjusting the remote focus only; seen as the in-focus image. No change in field size was seen on defocus by 100 µm. Panel 4E. Two fluorescent beads 5 µm diameter suspended in Sylgard resin, separated by 100 µm in depth, viewed simultaneously on the same camera. Upper: at primary focus, Lower: remote image;.

#### Magnification in the remote focus image

It has been reported that changes of magnification occur particularly at the edge of the field of view in scanning remote focus when the optical train is imprecisely set (Botcherby et al., 2013; Corbett et al., 2014) We tested the possibility that sub-optimal settings in the optical train imposed by the fixed microscope tube lens result in changes of magnification as the focus of the remote objective was adjusted. The focus of the direct and remote paths was set to be the same as seen in the oculars and the focus of O1 changed in 50 µm steps above or below, each time refocusing the remote image of the stage micrometer by translating O2 (see scheme in Fig. 2). The field diameter was measured on the stage micrometer in each z-position. The field diameter of 320 µm at zero displacement was unchanged with 100 µm z-displacement in remote focus as shown in Fig. 4C, D.

#### Tilted neuronal images

The ability to tilt neuronal images to maximise the dendritic area seen in epifluorescence is illustrated by Fig 6. Surviving cerebellar Purkinje neurons are usually oriented with dendrites deeper in the slice than the soma because of damage during slicing. To test the tilt, we used fixed preparations sparsely labelled with Gcamp6f (kindly provided by Drs J. Bradley and A. Jalil; mouse parasagittal cerebellum; age 30 days). In two example cells shown in Fig 6 the somatic and dendritic images (top pair A, B in each) are separated by 24 µm and 21 µm on the optical axis in the primary (ocular) image. They were brought into focus by adjusting the gimbal mirror on the remote focus path with 8.5° or 7.5 ° tilt, respectively, from primary focal plane.

**Fig. 5.**
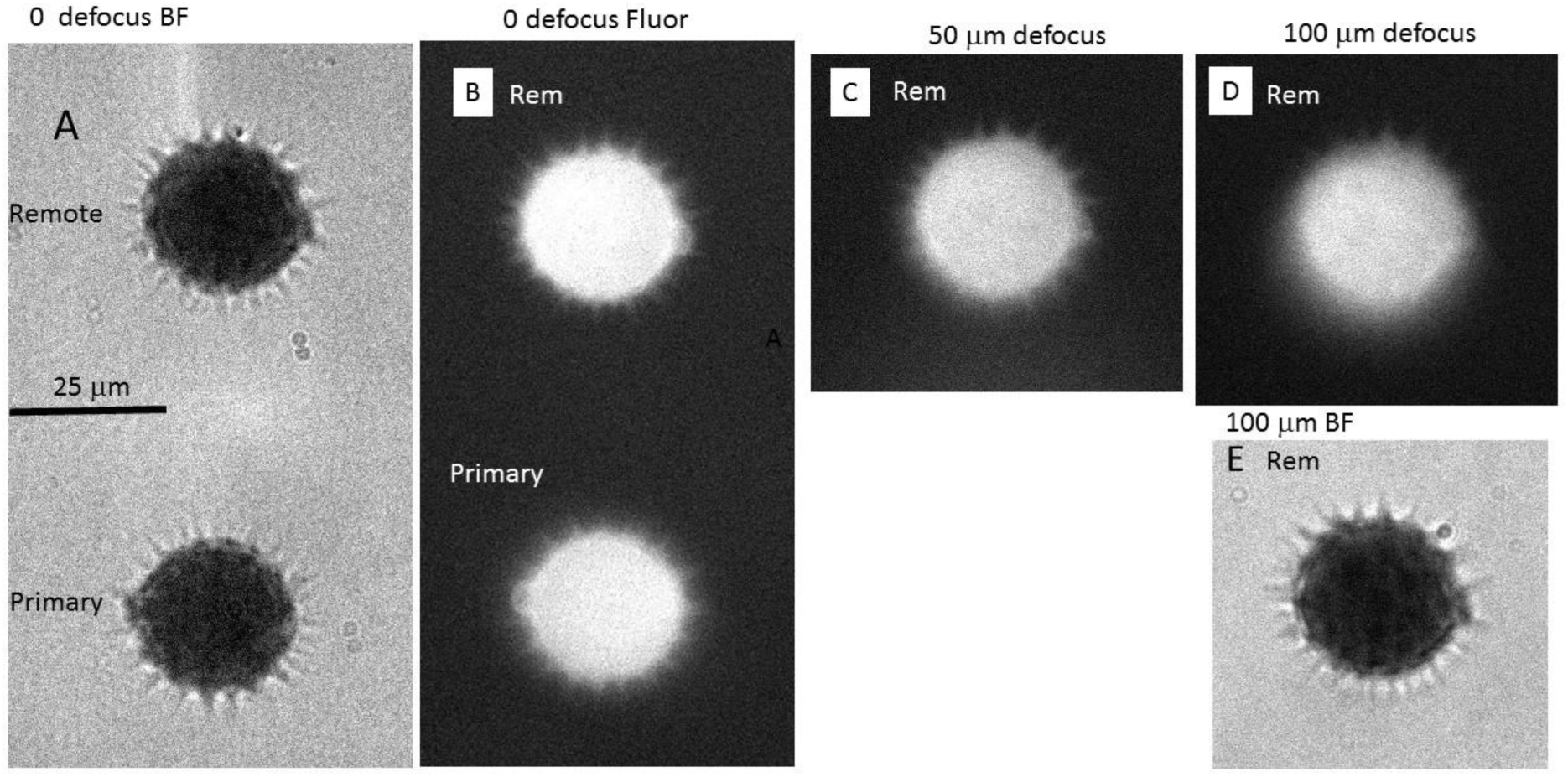
Comparison of resolution in the remote image at different depths below the primary focus. A: brightfield B: Fluorescence primary and remote images at the same depth. C: remote focus at 50 µm below primary; D, E: remote 100 µm below primary focus. Pollen grains 27 µm diameter imaged in brightfield or fluorescence (470 nm excitation, 530/40 nm emission). 500 ms exposures, EM gain 20. Calibration 25 µm.

**Fig 6.**
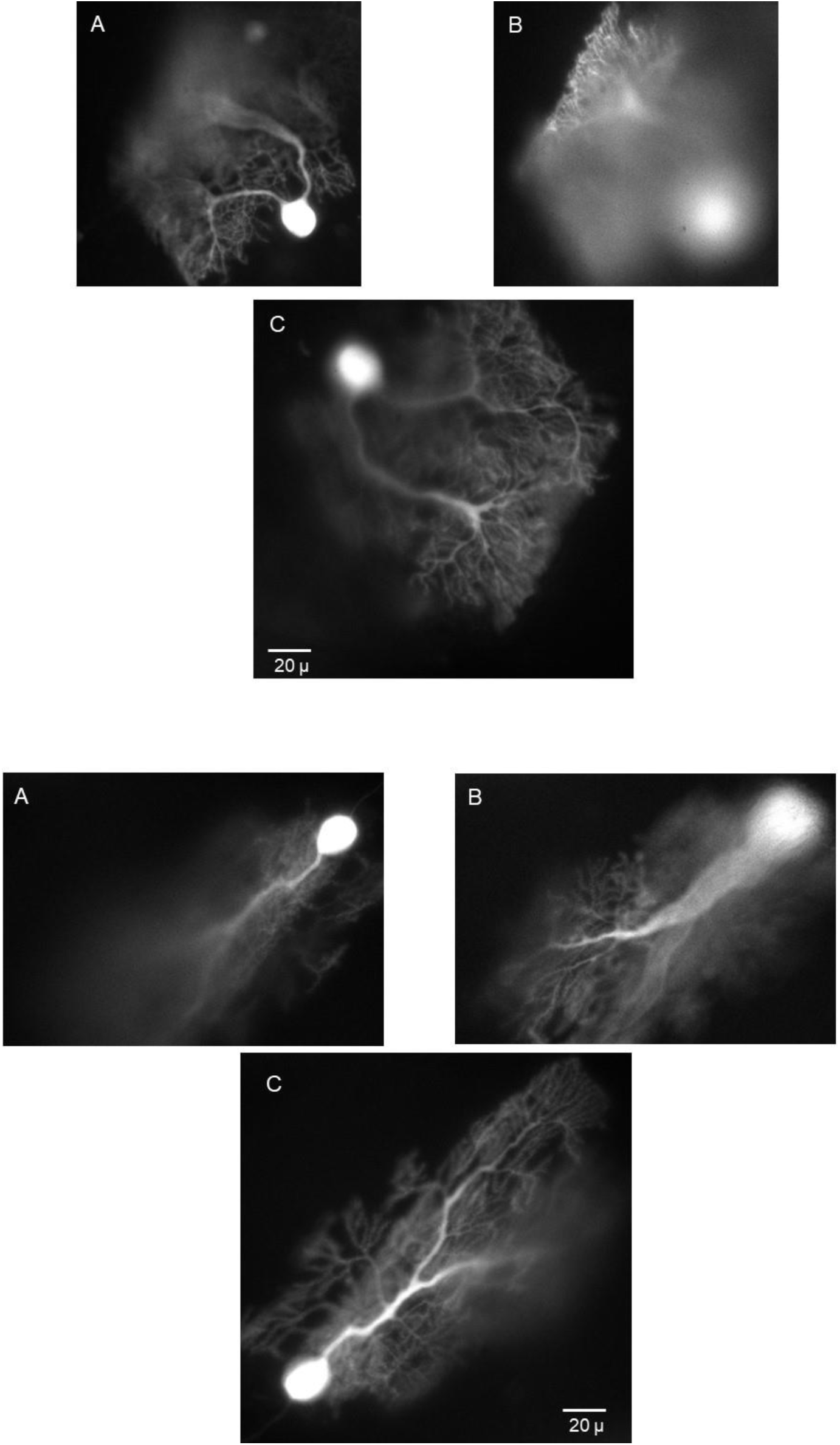
Tilting remote images. Two examples of Purkinje neurons of differing morphology are shown in (i upper) A-C and (ii lower) A-C. For both neurons Panels A show primary plane at the soma and (B) at the depth of the distal dendrites; panels (C) after adjusting remote mirror tilt to bring most of the dendritic tree into focus. The distal dendrites are 24 µm (i upper cell) and 21 µm (ii lower cell) below the soma. Note inversion in remote image. Field size in each image 180×180 µm. The pixel size (0.18 µm) does not resolve Purkinje neuron spines. Cells transfected with GCamP6f, mouse 30 days, parasagittal slice.

When imaging interneurons in a brain slice the somatic and axonal compartments are usually at different depths although intact within the slice. The ability to image the axon and dendrosomatic regions simultaneously permits activity in different regions to be correlated. Fig 7 shows images of a cerebellar Golgi interneuron tilted to maximise the view of the axon (left panel) and in the right panel a cerebellar molecular layer basket cell; both are tilted to bring most of the dendritic arbor and axon into the same image.

**Fig 7.**
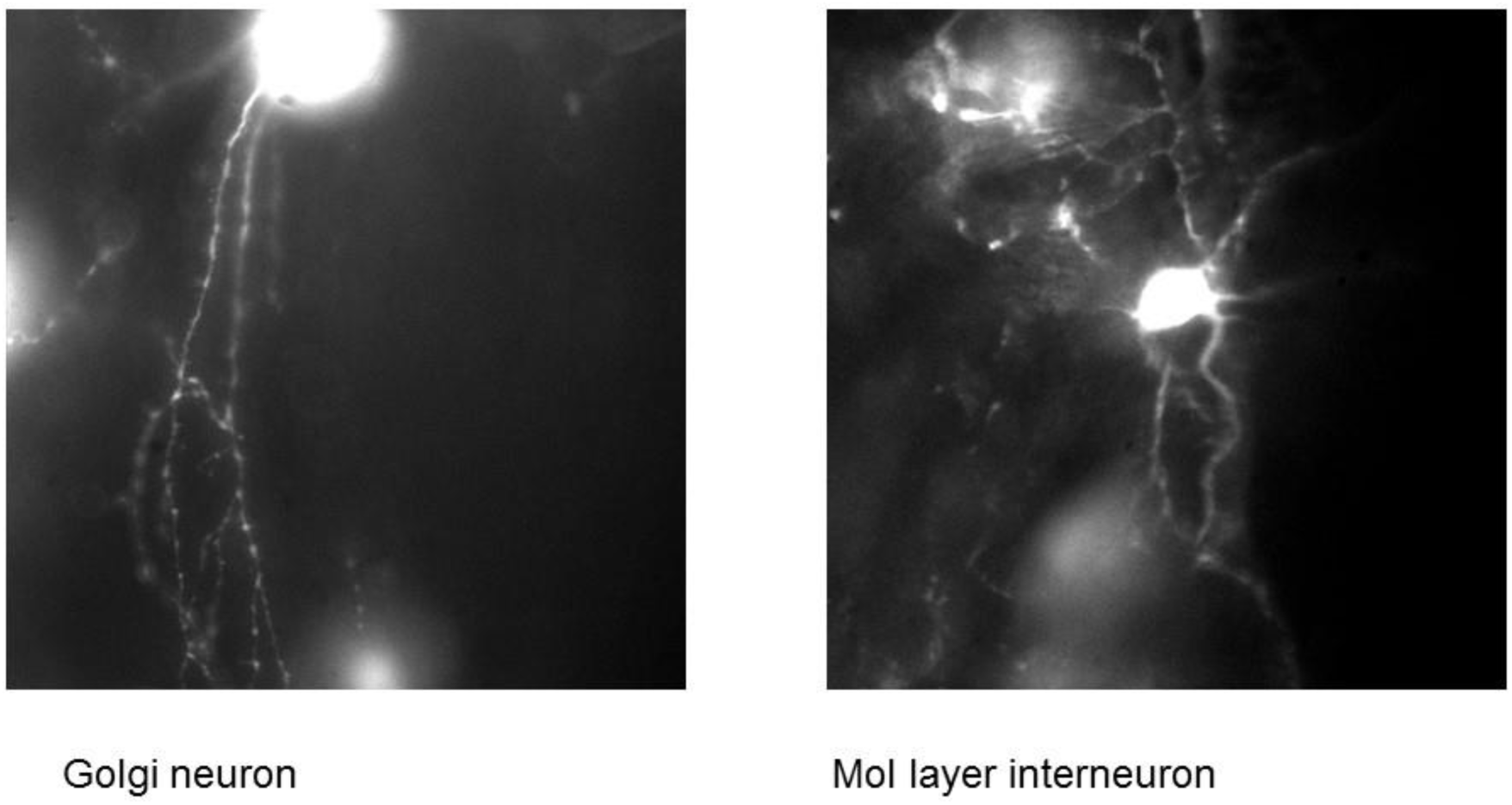
Images of a cerebellar Golgi cell (Left panel) and a molecular layer interneuron (Right panel) in sagittal slices with remote focus gimbal mirror adjusted to maximise the length of axon in focus in the tilted image. Cells sparsely labelled with eGFP. Field size 180 x 180 µm.

#### Imaging neurons at different depths in the slice

A larger field is desirable for viewing networks of neurons within the slice. To maximise the field a 40x 0.8 NA Leica water dipping was used at O1 and camera lens L2 changed to 100 mm focal length, projecting a larger field of 320 µm on the 12 mm camera sensor. This required adjustment of the optical train to have a tube lens L1 of 120 mm focus placed 120 mm from the intermediate image and adjustment of the position of components as discussed above. The fine position of L1 was adjusted to coincide the position and magnification of the image in the two arms.

Fig.8 shows images of interneuron networks obtained at different depths in sagittal cerebellar slices with the approaches described in Fig. 4. Each shows molecular layer interneurons labelled with GCamP6f at different depths viewed simultaneously. Fig 8A shows the image seen at the surface and Fig 8B the interneurons 30 µm deeper viewed with remote focus in the same location. The field size is 320 µm. Fig 8C shows a different region with the primary and remote images separated in a polarisation beam splitter and projected to adjacent regions of the same camera chip (scheme in Fig 4B; Cairn Optosplit), the upper image 35 µm deeper than the lower. Fig 8D shows images of molecular layer interneurons recorded at higher magnification, 180x 90 µm field, the upper part 50 µm deeper than the lower viewing the surface. The results show that in sparsely labelled neural preparations simultaneous fluorescence imaging at two depths in a slice is feasible with remote focus.

**Fig 8.**
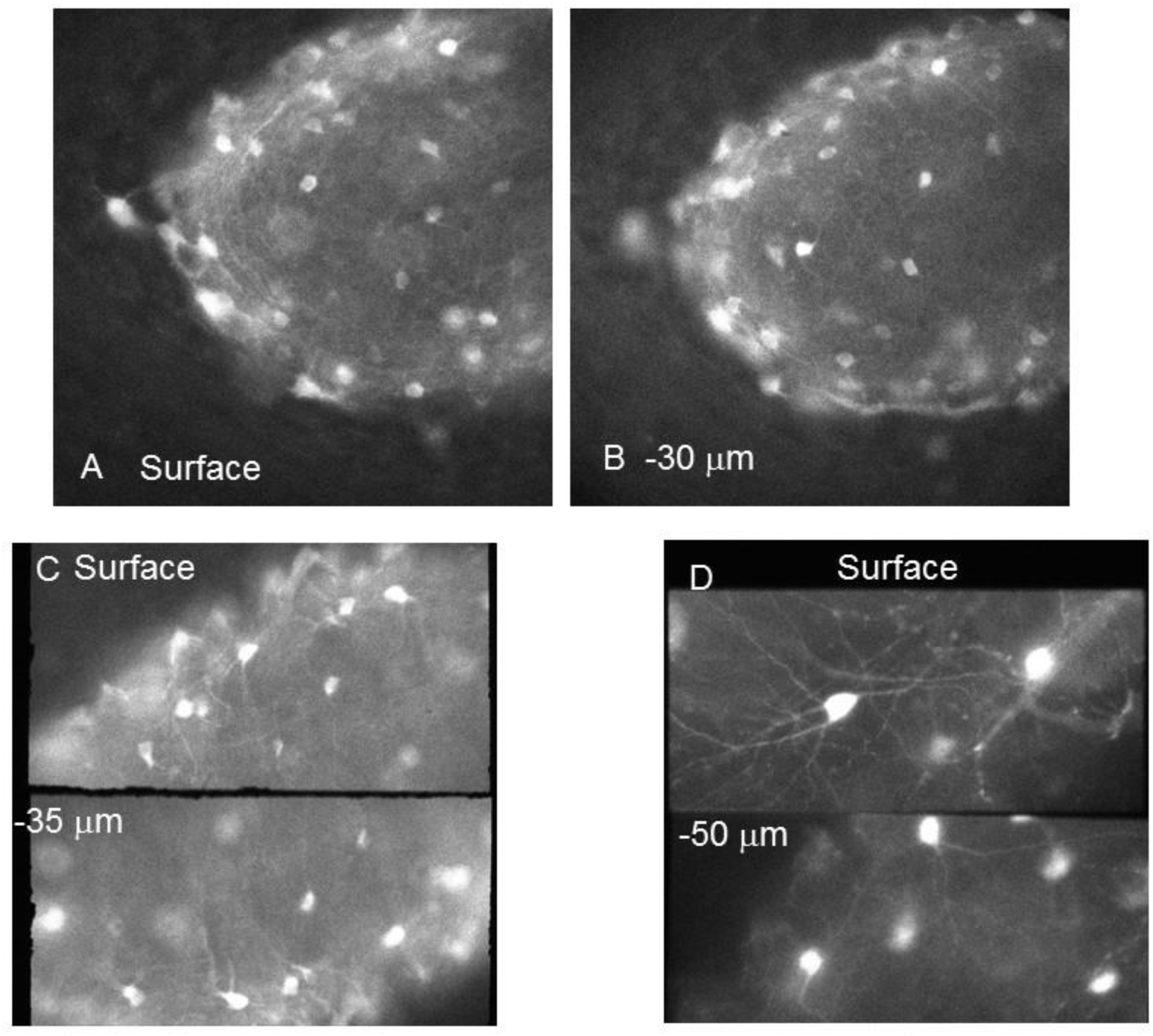
Simultaneous imaging at 2 depths. Cerebellar molecular layer interneurons imaged in 2 sagittal planes. Fig.8A. Surface of a slice with molecular layer interneurons expressing GCam6f. Field size 320 x 320 µm. 8B. view in the remote path 30 µm deeper in the slice (remote focus image inverted for display). Panel C. Interneurons in 2 sagittal planes separated by polarization in an Optosplit 2 (Cairn Research) onto the same camera sensor. Upper: at the surface, Lower: remote focus image 35 µm deeper. Note optical inversion of remote image (uncorrected). Panel D. Higher magnification images of interneurons (180 µm field width) separated by polarization at 2 depths 50 µm apart. Lower pane: slice surface; Upper pane: 50 µm deeper (Image inversion uncorrected). Mouse, 4 months, cerebellum, sagittal slices, Gcam6f positive neurons. 470/40 nm excitation, 530/40 nm emission. Each full panel is 1000×1000 pixels, Optosplit panels C. 320×160 µm and D 180×90 µm, 70 ms exposure.

##### Time-resolved imaging at 2 depths

The ability to image with fast acquisition simultaneously at 2 depths was investigated with zebrafish embryos selectively expressing GCamP5 in motor neurons of the spinal cord in the *Tg(mnx1:GCamP5G)* transgenic line. Spontaneous activity in motoneurons and axons was recorded as fluorescence changes in GCamP5 at 24-30 hours post-fertilisation. Muscle excitation was blocked with alpha-bungarotoxin to prevent contraction. The fluorescence changes due to activation of GCamP5 by Ca influx were recorded at 2 depths in motoneuron chains from rostral and caudal regions of 9 fish. The layers of neuronal soma were separated by 30-40 µm across the midline, in the example shown 34 µm, separation measured in each case by refocusing between primary and remote images. The activity in a rostral region of the tail of a fish at 28 hours post-fertilisation is shown for contralateral sets of neurons in Fig. 9. Full frame images of 180 microns were split into 2 polarisations and projected onto adjacent regions of the camera sensor (180×90 microns, Andor Ixon 887 2×2 binned giving 0.8 x 0.8 µm pixels; Multisplit, Cairn Research.; Fig.9A). Regions of interest in the output axons innervating myotomal muscle were selected and the time course plotted for 2000 frames at 30 Hz (Fig. 9B). Increases of GCamP5 fluorescence due to integrated bursts of spikes in the axons and soma showed reciprocal activity between neurons in the two regions separated here by depth, as reported previously (Warp et al 2012, Muto et al 2011). Similar results were obtained in 18 recordings from 9 zebrafish. Although the kinetic response of GCamP5 is too slow to resolve single spikes, alternate bursts are detected from the two opposing lateral networks with good signal/noise in both polarisations, corresponding to direct and remote focus images at different depths in the preparation. Although reciprocal activity in neural soma of opposite sides has been shown in dorsal views of neuron soma at the same focus (Muto et al., 2011, Warp et al., 2012) here we additionally obtain data from axonal connexions to the myotome in two dorso-ventral planes imaged simultaneously.

**Fig. 9.**
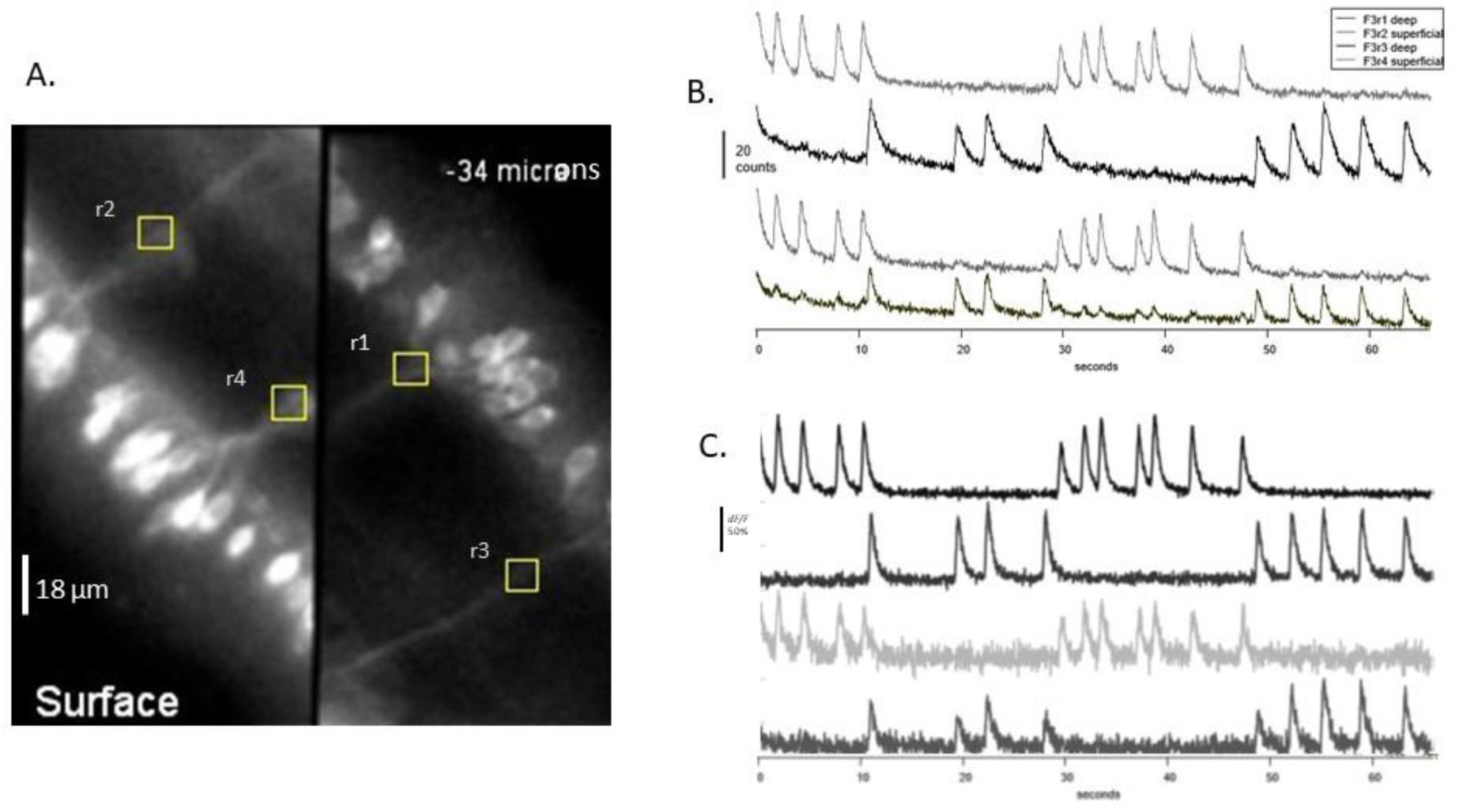
Time resolved fluorescence in spinal neurons of Zebra fish recorded simultaneously at 2 depths. Soma of labelled motoneurons separated by 34 µm across the midline in the spinal cord in transgenic *Tg(mnx1:GCamP5G)* zebrafish embryos mounted horizontal in agar. Motoneurons expressed GCamP5, spontaneous activity in motor neurons was recorded 28 hours post-fertilisation at 23° C. Muscle excitation was blocked with alpha-bungarotoxin. Panel A: Left part shows neurons situated beneath the uppermost lateral surface, rostral aspect to the top. Right part shows neuronal cell bodies 34 µm deeper, caudal aspect top because of image inversion in the remote objective. Images separated by polarisation and projected in a Multisplit (Cairn Research; a polarising cube beamsplitter replaces first dichroic reflector) to adjacent regions of the sensor of Andor Ixon 887 EMCCD. Regions of interest r1-r4 12×12 pixels defined as shown. Excitation 470/40 nm, Emission 530/40 nm. Panel A Sum of 10 images to show morphology and ROI. Panel B: Fluorescence, raw data;1000 frames acquired at 17 ms exposure, mean values from each of 4 regions indicated in A are plotted with time. Frame rate 30 Hz, EM gain 100, 2×2 binning, each image 180 x 90 µm. Panel C: Raw data converted to *dF/F*_*0*_ after background subtraction and high pass spatial filter; *F*_*0*_ is the average of 5 frames prior to the fluorescence transient,.

## Discussion

Camera based imaging of fluorescent indicators combined with electrophysiological recording is widely used in neuroscience to correlate neuronal output with localised ion concentration or local membrane potential changes. Camera-based imaging of emission from fluorescent indicators has two advantages over laser scanning methods, the simultaneous acquisition of data over the field of view and high time resolution achievable with recent advances in camera technology. In conventional wide-field imaging a major technical difficulty is in monitoring fluorescence over neuronal structures that do not lie in the conventional focal plane, requiring refocus and precluding simultaneous multisite analysis. As shown previously, remote focus in a second objective permits fluorescence recording in tilted non-normal focal planes within the remote image (Anselmi et al, 2011). There are no reports of its implementation in the emission path of camera-based systems in conjunction with electrophysiological recording. Descriptions of remote focus applied to excitation in laser scanning microscopes in conjunction with electrical or mechanical measurements have been given previously but differ substantially in implementation from camera based fluorescence imaging (Botcherby et al., 2012; 2013; Sofroniew et al., 2016, Zhuan et al.,2019). Here we have adapted remote focus to fit with the practical constraints of electrophysiological microscopes. We have made use of the non-remote polarisation to simultaneously view the conventional ocular image, providing a reference image and an aid to alignment as well as a second imaging depth. We describe image tilt applied to Purkinje neurons and cerebellar interneurons in tissue slices, we show simultaneous imaging of neuron fields at 2 well separated depths in the slice that can be tilted with respect to the conventional focus.

The remote focus method was described in several formats and evaluated first by Botcherby and coworkers (2008). It has been shown to be robust with regard to resolution over the range of depths needed in slice recording, and relatively tolerant of imperfect alignment of the optical train by Anselmi etal (2012) Qui et al., (2014) and here. Botcherby et al., (2008) found a range of +/− 80 µm defocus in the remote image from the primary image before aberrations were seen (1.4 NA oil primary objective). In conditions similar to those here (40x, 0.8 NA, water) Anselmi et al., 2011 found a range of +/− 150 µm. No change in the Strehl ratio was found by wavefront analysis over this range and the resolution was insensitive to misalignment of the pupils of primary and remote objectives. One potential problem in commercial microscopes is the fixed distance from objective to the tube lens usually closer than is optimal for remote imaging. However, the images obtained indicate that spatial resolution nevertheless remains good up to 150 µm depth even in the head focus Zeiss microscope used here (tube lens 165 mm focal length, distance to back pupil 110mm); in other physiological microscopes (Olympus BXF or Scientifica Slicescope equipped with epifluorescence condenser) the tube lens-pupil distance is closer to the optimum for a 4f relay of the image. It may also be noted that the cone angle of the remote objective here is greater than either of the water dipping objectives 63x or 40x used at O1 and is not expected to impair resolution (Botcherby et al., 2008).

For time resolved imaging large, 1 µm binned pixels are often used in kinetic measurements at high frame rates, and loss of spatial resolution is less of a concern compared with high resolution imaging applications. The loss of photons in the remote focus is potentially more important, the measurements indicate that 30% of light at the intermediate image is transmitted in one polarisation *via* the remote optics to the camera. Measurements from both remote paths combined at the camera gave 65% throughput from the intermediate image and could be employed to increase light throughput. In thick biological preparations scattering of emitted photons within the tissue will also contribute to reduced resolution; however this was not found to be a problem in the sparsely labelled tissues used here.

Imaging simultaneously at two widely separated depths in the preparation as described here is novel and may be particularly useful when communication between tissue layers is physiologically important, such as in the retina, or where activity in adjacent layers may give information about the inputs and outputs in adjacent circuits, for example as in sagittal circuits of the cerebellar cortex. Further, image splitting is done with polarisation in the first pass to separate the depths, and each path can be further split to two colours of fluorescence at each level with no additional light loss.

In time-resolved imaging of GCamP5 from spinal neurons at two depths in Zebrafish tail (Fig 9) there was contamination of fluorescence between the two imaging planes when large ROIs were used. Because the optical manipulations of remote focus are in the emission path, confocal methods that use structured illumination or speckle in the excitation with subsequent HiLo image processing in emission (Lauterbach et al., 2015; Ventalon and Mertz, 2005, 2006, Lim et al., 2011) may be used to remove out of focus fluorescence. The z-projection inherent in laser speckle would be useful in the present context at 2 imaging depths.

## Acknowledgements

We thank Ali Jalil and Jonathan Bradley for kindly providing sparsely labelled GCamP6 or eGFP labelled fixed tissues, Marcel Lauterbach for helpful comments on the manuscript and discussion.

